# Pisces: An Accurate and Versatile Variant Caller for Somatic and Germline Next-Generation Sequencing Data

**DOI:** 10.1101/291641

**Authors:** Tamsen Dunn, Gwenn Berry, Dorothea Emig-Agius, Yu Jiang, Serena Lei, Anita Iyer, Nitin Udar, Han-Yu Chuang, Jeff Hegarty, Michael Dickover, Brandy Klotzle, Justin Robbins, Marina Bibikova, Marc Peeters, Michael Strömberg

## Abstract

**Motivation:** Next-Generation Sequencing (NGS) technology is transitioning quickly from research labs to clinical settings. The diagnosis and treatment selection for many acquired and autosomal conditions necessitate a method for accurately detecting somatic and germline variants, suitable for the clinic.

**Results:** We have developed Pisces, a rapid, versatile and accurate small variant calling suite designed for somatic and germline amplicon sequencing applications. Pisces accuracy is achieved by four distinct modules, the Pisces Read Stitcher, Pisces Variant Caller, the Pisces Variant Quality Recalibrator, and the Pisces Variant Phaser. Each module incorporates a number of novel algorithmic strategies aimed at reducing noise or increasing the likelihood of detecting a true variant.

**Availability:** Pisces is distributed under an open source license and can be downloaded from https://github.com/Illumina/Pisces. Pisces is available on the BaseSpace™ SequenceHub as part of the TruSeq Amplicon workflow and the Illumina Ampliseq Workflow. Pisces is distributed on Illumina sequencing platforms such as the MiSeq™, and is included in the Praxis™ Extended RAS Panel test which was recently approved by the FDA for the detection of multiple RAS gene mutations.

**Supplementary information:** Supplementary data are available online.

## 1 Introduction

Next-Generation Sequencing (NGS) technology is transitioning quickly from research labs to clinical settings. The diagnosis and treatment selection for many acquired and autosomal conditions necessitate a method for accurately detecting somatic and germline variants (Dietel et al., 2015; Dong et al., 2015). Many algorithms have been developed for somatic single nucleotide variant (SNV) detection in matched tumor-normal DNA sequencing, and many algorithms have been developed for detecting germline variants, GATK being the most well-known amongst them (DePristo et al., 2011; McKenna et al., 2010). However, there is no single front runner, and different callers dominate in different situations. (Dietal et al, 2006; Horak et al., 2016; Sandmann et al., 2017; Xu et al., 2014). Particularly in the context of amplicon workflows, the standardization of variant calling pipelines remains elusive. (Betge et al., 2015).

Pisces is uniquely designed to perform well on germline and somatic amplicon data, particularly in the common situation where no matched normal sample exists for a given tumor sample. Pisces Suite is a set of simple-to-use command-line applications that are easily incorporated into more complex workflows. Pisces requires only aligned sequence data and related reference genome files as input. Pisces returns a variant call file (VCF) with SNVs and small insertions, deletions, and (optionally) complex variants such as phased small variant recompositions. Pisces is restricted to small variant calling (it does not call structural variants), and is best suited to amplicon sequencing. However, Pisces is particularly versatile in the sense that it outperforms many other variant callers in both the germline and somatic contexts, and also in the modularity of its elective architecture. Here we present an overview of the Pisces Suite. We compare Pisces performance with a number of alternative small-variant calling tools and give a summary of the performance results for our set of samples.

## 2 Methods

Pisces execution is organized into four sequential modules, which may be invoked as needed depending on the type of sample to be processed. Each module incorporates a number of unique algorithmic strategies aimed at reducing noise or increasing the likelihood of detecting a true variant, which we summarize in the next section. We show how each individual module successively contributes to enhanced accuracy in the SupplmentaryResults. All but the core variant calling step may be omitted if the sample does not necessitate additional processing. (For example, an FFPE treated somatic sample would be a good candidate for the full suite, while a fresh-frozen germline sample could be accurately called with only the small variant calling step.) This versatility, as well as its accuracy, make the Pisces Suite a favorable tool for use in small variant detection.

### 2.1 Pisces Overview

The Pisces Suite workflow comprises four sequentially executed modules:

1. Pisces Read Stitcher: Reduces noise by stitching paired reads in a BAM (read one and read two of the same molecule) into consensus reads. The output is a stitched BAM.
2. Pisces Variant Caller: Calls small SNVs, insertions and deletions. Pisces includes a variant-collapsing algorithm to coalesce variants broken up by read boundaries, basic filtering algorithms, and a simple Poisson-based variant confidence-scoring algorithm. The output is a VCF.
3. Pisces Variant Quality Recalibrator (VQR): In the event that the variant calls overwhelmingly follow a pattern associated with thermal damage or FFPE deamination, the VQR step will downgrade the variant Q score of the suspect variant calls. The output is an adjusted VCF.
4. Pisces Variant Phaser (Scylla): Uses a read-backed greedy clustering method to assemble small variants into complex alleles from clonal subpopulations. This allows for the more accurate determination of functional consequence by downstream tools. The output is an adjusted VCF.

More information on the Pisces algorithms are given in the Supplementary Methods section.

### 2.2 Implementation and Performance

The Pisces Suite is implemented in the C# programming language and can be run on multiple platforms using the .Net core technology. Multithreaded options are available for all time-consuming modules. Pisces processing speed scales well with BAM size: An average runtime for the Pisces Variant Caller on a 470 MB BAM (8 million reads) is 85 seconds. Runtime for a 2 GB BAM (60 million reads) is about 4 minutes, and for a 65 GB BAM (900 million reads), about 40 minutes (35 seconds per GB processed). These examples were run with 20 threads, on a Linux Cluster node with Intel(R) Xeon(R) CPU E5-2670 @ 2.60GHz processors.

### 2.3 Testing Methodology

We compared Pisces performance with a number of alternative small-variant calling tools. The selection of third-party tools was based on the principle that they showed a superior performance in previous benchmarking studies (either somatic or germline) (Dietal et al, 2015; Horak et al., 2016; Sandmann et al., 2017; Xu et al., 2014). Each tool chosen offers a slightly different variant calling strategy, and might be optimal in other situations. We do not aim to give a comprehensive comparison of small-variant calling tools as this is given elsewhere (Sandmann et al.,2017; Xu et al., 2014). What follows is an overview of our test-generation strategy and a summary of the performance results for our set of samples.

To demonstrate the accuracy of each variant caller, we generated BAMS from four amplicon datasets (two for the somatic workflow and two for the germline workflow) where truth was known. The germline datasets (the Variant Panel Dataset and the Myeloid Panel Dataset) were generated using cell line samples, and the somatic datasets (the Titration Dataset and the RAS Panel Dataset) were generated from a titration mixture and a set of FFPE-treated solid tumor samples of unknown heterogeneity. The selection of the datasets was motivated by a desire to test the variant callers on well-characterized samples, and also on realistic cancer samples that were outside any common training set.

### 2.4 Germline Datasets

All germline testing was done using established cell line samples from individuals NA12878 and NA12877 from the Coriell Institute, New Jersey. High-confidence variant calls are available for these individuals via Platinum Genomes build 2016-1.0 (Eberle et al., 2017). These accepted variant calls form a gold standard for evaluating germline calling. The two germline datasets (The Variant Panel Dataset and Myeloid Panel Dataset) were created using the same established cell lines, but different panels.

The **Variant Panel** was designed to target known variants in the NA12878 and NA12877 samples, specifically for the purpose of assessing the accuracy of sequencing applications. It includes 91 high-confidence SNVs and 42 high-confidence indels identified by Platinum Genomes. The variant panel samples correspond to the “VP” dataset labels in Table 1.

**Table 1.**
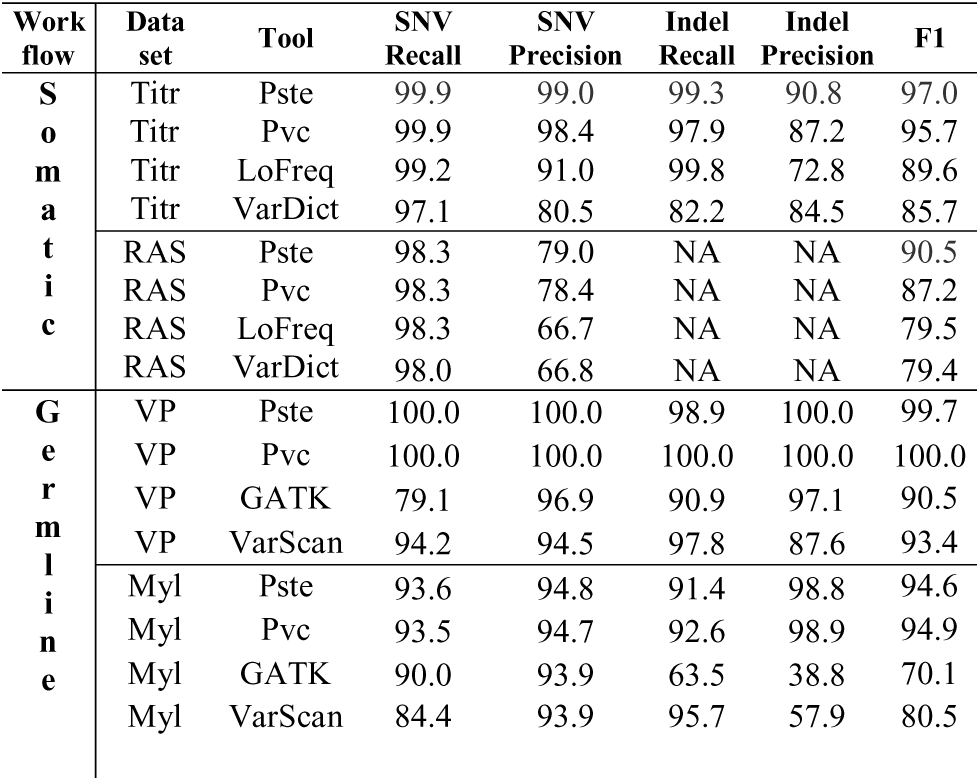
We show average accuracy metrics by variant caller across all samples in each of the four datasets. The single F-score (F1) given is the average of the F1 for SNVs and the F1 for indels. Pste means the full Pisces Suite was used (specifically, the stitcher, variant caller and the recalibration step) and Pvc means only the pisces variant caller was used. The phasing step (Scylla) was deliberately omitted here, since Hap.py cannot be used to assess the accuracy of complex allele reconstruction. Scylla results are discussed in the Supplementary Results section.

The **Myeloid Panel** is a commercially available panel (Illumina TruSight^®^ Myeloid Sequencing Panel) and specifically targets genes frequently mutated in blood cancer disorders. Of those targeted variants, 84 are high-confidence variants identified by Platinum Genomes. The myeloid panel samples correspond to the “Myl” dataset labels in Table 1.

### 2.5 Somatic Datasets

The somatic datasets (the Titration Dataset and the RAS Panel Dataset) include one engineered dataset and one clinical dataset typical to oncology research.

The **Titration** dataset was built to extend the accuracy assessment capabilities of the Variant Panel into the somatic context. The Titration dataset is a mixture of the NA12878 and NA12877 cell line samples described above, serially diluted to 16%, 12%, or 8% by volume. Since the datasets include heterozygous variants, and due to sequencing variability, many confirmed variants were only observed at much lower frequencies, down to 1%. The titrated samples were run with the Variant Panel, and therefore cover the same high-confidence SNVs and indels identified by Platinum Genomes. The titration samples correspond to the “Titr” dataset labels in Table 1.

The **RAS Panel** dataset was generated from a set of colorectal cancer tissue blocks which were formalin-fixed paraffin-embedded (FFPE) treated on or before 2005, and extracted eight to nine years later. Those same biological samples were additionally evaluated by one of two orthogonal methods to provide gold standard variant calls. The orthogonal methods used were Sanger sequencing and therascreen *KRAS* test (Qiagen). The RAS Panel samples correspond to the “RAS” dataset labels in Table 1.

## 3 Results

### 3.1 Evaluation Strategy

We generated four sets of BAM files corresponding to the four datasets described previously. The BAM files were generated using the Illumina amplicon aligner, an implementation of the Smith-Waterman-Gotoh algorithm (Gotoh, 1982). Each set of BAM files was processed through the variant caller, and the resulting VCFs were assessed using the Hap.py accuracy assessment tool (https://github.com/Illumina/Hap.py).

The variant callers used for the analysis are the GATK HaplotypeCaller (McKenna et al., 2010), LoFreq (Wilm et al., 2012), Pisces (https://github.com/Illumina/Pisces), VarDict (Lai et al., 2016) and VarScan (Koboldt et al., 2013). Precision and recall were assessed using Hap.py. The input data is freely available, and links are included in the supplementary results section.

### 3.2 Comparison Results

We see that all variant callers performance is often very similar for SNV recall, but indel precision varies dramatically. We also see that the lowest performance (from all callers) is on the RAS panel dataset. This underscores the importance of including real cancer sample datasets when assessing variant calling capabilities. In each of the four datasets, Pisces attained the largest number of best-performing metrics amongst the tools used. For germline calling, Pisces Variant Caller alone does just as well (slightly better) than the more complex pipeline. However, in the somatic case, best results are achieved with the full Pisces Suite. We discuss the accuracy gains (and losses) attributable to each optional Pisces Suite module engaged in more detail in the Supplementary Results section. To conclude, Pisces is a versatile and accurate tool for small-variant detection on amplicon datasets.

## 4 Acknowledgements

The GATK3 was made available for use by Illumina through the generosity of the Broad Institute. VarScan, VarDict, LoFreq are all freely available on GitHub.com. The Pisces team is profoundly grateful to the authors’ generosity and commitment to open source and open science.

